# Exposure to PFOA and GenX elicits cell type-specific impacts on p53 and TGF-β signaling pathways

**DOI:** 10.1101/2025.05.03.652004

**Authors:** Hongran Ding, Maya Slack, Holly McClure, Wen Gu, Helene Gu, Ferdinand Kappes, Thomas F. Schultz, Jason A. Somarelli, Anastasia Tsigkou

## Abstract

Per- and polyfluoroalkyl substances (PFAS) are a class of synthetic chemicals extensively used as plastic additives. Their environmental persistence and potential for bioaccumulation have raised significant toxicological concerns. This study evaluates the cytotoxic and molecular effects of Perfluorooctanoic acid (PFOA) and its replacement compound, Perfluoro(2-methyl-3-oxohexanoic) acid (GenX), in human-derived skin (A375), liver (HepG2), kidney (SN12C), and colon (SW620) cell lines. The experimental design assessed cell viability, gene expression, and perturbations in key cellular stress pathways, with a particular focus on TGF-β/SMAD-mediated inflammation and the p53-driven DNA damage response.

Our results demonstrate compound- and cell-type-specific toxicity, with GenX displaying reduced cytotoxicity compared to PFOA across all cell types. Molecular analyses revealed that both PFAS compounds induced alterations in the TGF-β/SMAD pathway, consistent with a pro-inflammatory cellular state. Additionally, we observed activation of the DNA damage response, as evidenced by increased expression of ATM, ATR, and p53, alongside ribosomal stress-related changes in RPL5 and RPL11. Notably, while skin and liver cells exhibited similar response profiles, kidney and colon cells showed divergent modulation of SMAD signaling, suggesting tissue-specific susceptibility and mechanistic differences.

These findings contribute to a deeper understanding of the differential toxicological profiles of legacy and replacement PFAS, with implications for health risk assessment and regulatory policy.

**Graphical Abstract:** 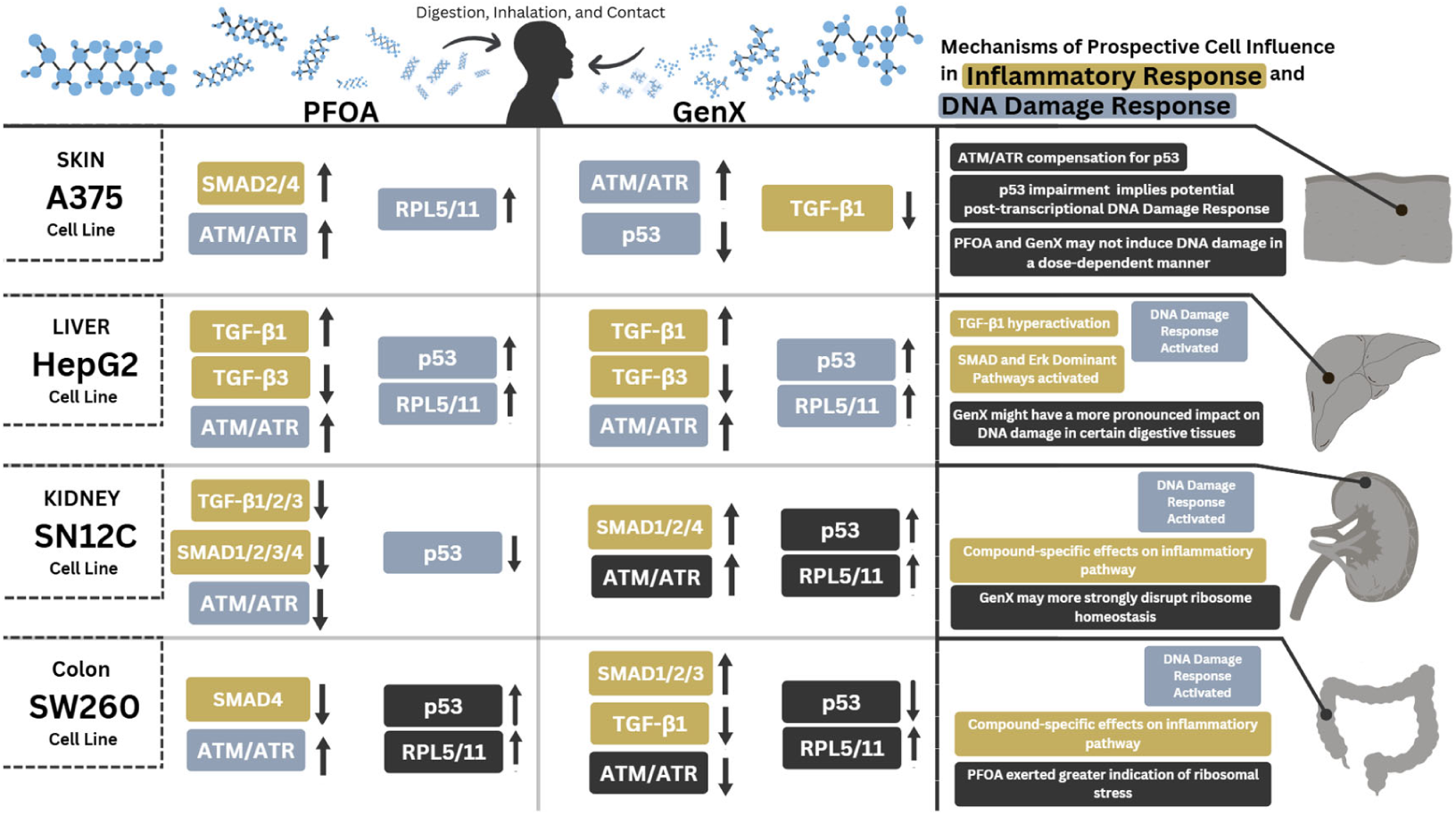

## Introduction

Per- and polyfluoroalkyl substances (PFAS) are a large class of synthetic fluorinated compounds designed for their water and oil repellent properties, making them widely used as additives in plastics and consumer products such as cosmetics, food packaging, pesticides, textiles, and medical devices.^1,2,3^ However, during manufacturing disposal, PFAS are released to the environment and accumulate in plants.^4,5^ Their highly stable carbon-fluorine bonds make them exceptionally resistant to degradation, leading to persistence in soil, water, and air, as well as bioaccumulation in organisms.^1,6^

Human biomonitoring confirms widespread exposure. For example, PFAS have been detected in biospecimens from Guangdong Province, China, with mean concentrations of 3.6 ng*mL^-1^ and 5.4 ng*g^-1^, of PFOA in hair and nails, respectively.^7^ Similarly, dietary intake studies in Norway revealed exposure to 18 PFAS compounds and 12 precursors, with perfluorooctane sulfonate (PFOS) and PFOA as predominant serum contaminants.^8^ Environmental surveys further demonstrate contamination: PFOA concentrations range from 1.49 ng/g - 8.0 ng/g in soil^3,9^ and 0.1 ng/L - 3.9 ng/L in groundwater, with highly contaminated regions such as the Yangtze River in China reaching median PFOA levels of 297.5 ng/L.^10^ While most toxicological studies focus on serum levels or liver toxicity, recent evidence indicates that PFAS, including legacy compounds like PFOA and emerging replacements such as GenX, accumulate differentially across organs—liver, kidney, lung, colon, and bone—suggesting that tissue-specific responses are likely and under-characterized.^11, 12, 13^ These differences in distribution highlight the need to investigate organ-level mechanistic responses to understand the full spectrum of PFAS toxicity.

### Health Impacts of PFOA and Emergence of GenX

Among more than 10,000 PFAS, perfluorooctanoic acid (PFOA) is one of the most common, well-known, and extensively studied chemicals in this class. PFOA exposure has been associated with a wide range of adverse health effects, including immunotoxicity,^11^ endocrine disruption, dyslipidemia, and metabolic abnormalities, such as hypercholesterolemia and hyperuricemia.^14,15^ In animal models, PFOA accumulates in the liver, leading to hepatocellular hypertrophy and other toxic effects.^16^ Due to its persistence and harmful health associations, PFOA was added to the Stockholm Convention on Persistent Organic Pollutants in 2019, resulting in near-global restrictions.^17^

As a replacement, GenX (hexafluoropropylene oxide dimer acid) was introduced in the early 2000s.It was marketed as a safer alternative due to its shorter chain length and ether group conferring a reduced environmental half-life.^18^ However, emerging evidence suggests that GenX may also be toxic. Animal studies have reported hepatotoxicity,^16^ peroxisome proliferator-activated receptor (PPAR) pathway activation with downstream effects on lipid metabolism,^19^ induction of apoptosis in hepatocytes, and possible epigenetic toxicity.^20^ Increasingly, GenX has been described as a “regrettable substitute”.^18^ While human population data for GenX remain limited, importantly, human biomonitoring data have begun to emerge: serum and urine analyses in exposed U.S. communities confirm measurable GenX levels, indicating direct human uptake and systemic exposure.^21,22^ Based on available information, the EPA has marked the liver to be sensitive from oral exposure to GenX.^23^

While most PFAS studies have focused on serum burden or liver effects, recent findings indicate tissue-specific accumulation in kidney, lung, and bone.^11^ A 2024 EPA research project tested PFAS across eight subclasses using six human cell types, revealing cell-type specific activities, but the extent to of this activity is little explored.^24^ Mechanistic work is beginning to show that PFAS perturb cellular stress signaling. In particular, PFOA has been linked to transforming growth factor-β (TGF-β) signaling, a central pathway in fibrosis, inflammation, and tissue homeostasis.^25^ Elsheikh et al^26^ demonstrated that oral PFOA exposure in rats induced oxidative stress, pulmonary fibrosis, and upregulation of TGF-β1, Smad2, and Smad3, alongside Smad7 downregulation. Notably, withdrawal of PFOA partially reversed these effects, highlighting the dynamic nature of this pathway. Similarly, GenX has been associated with TGF-β activation and intestinal barrier disruption.^27^

Furthermore, other critical stress pathways remain underexplored in the context of PFAS exposure. Preliminary studies suggest that PFAS may perturb DNA damage responses.^28^ Central to this process are the tumor suppressor p53 and its upstream regulators ATM and ATR, which orchestrate DNA repair, apoptosis, and cell cycle arrest.^29^ Although some reports have linked PFAS to DNA damage and oxidative stress,^28,30^ the evidence remains sparse and fragmented. Similarly, ribosomal stress mediators RPL5 and RPL11, which stabilize p53 under nucleolar stress,^31^ represent novel and largely unexplored PFAS targets. To date, there is little to no direct evidence on PFAS-mediated regulation of ribosomal stress, highlighting an important gap in our understanding of how these compounds affect cellular homeostasis beyond canonical signaling pathways. Taken together, these gaps underscore the need for comprehensive studies that directly compare the effects of legacy and replacement PFAS across multiple tissues and cell types. Existing evidence has suggested limited associations between PFAS exposure and regulators such as ATM, ATR, and SMAD4,^32,33^ yet it is unclear whether these interactions are consistent across organ systems or differ between long- and short-chain compounds. By incorporating novel targets such as RPL5 and RPL11, it becomes possible to investigate whether ribosomal stress functions as an additional focal point for PFAS toxicity. Comparative, cross-tissue analyses are therefore essential to determine if legacy compounds such as PFOA and replacement chemicals such as GenX elicit distinct mechanistic responses, or whether they converge on common stress pathways with tissue-specific variations. Our study addresses this gap by systematically interrogating these pathways, thereby advancing the understanding of PFAS-induced cellular stress and potential carcinogenic risk,^33^ yet it is unclear whether these interactions are consistent across organ systems or differ between long- and short-chain compounds. By incorporating novel targets such as RPL5 and RPL11, it becomes possible to investigate whether ribosomal stress functions as an additional focal point for PFAS toxicity. Comparative, cross-tissue analyses are therefore essential to determine if legacy compounds such as PFOA and replacement chemicals such as GenX elicit distinct mechanistic responses, or whether they converge on common stress pathways with tissue-specific variations. Our study addresses this gap by systematically interrogating these pathways, thereby advancing the understanding of PFAS-induced cellular stress and potential carcinogenic risk.

### PFAS and Cellular Stress Pathways

Although PFAS are often characterized in terms of metabolic and immunotoxic outcomes, recent studies suggest they also perturb cellular stress signaling pathways. As previously mentioned, PFOA has been shown to modulate the transforming growth factor-β (TGF-β) pathway, which regulates fibrosis, inflammation, and tissue homeostasis.^26^ Importantly, some effects were reversible following withdrawal, suggesting dynamic regulation of this pathway. Similarly, GenX has been associated with activation of TGF-β signaling in the liver and intestine, alongside epithelial barrier disruption.^27^

### Knowledge Gaps and Rationale

While evidence indicates that both legacy PFAS (PFOA) and replacements (GenX) can disrupt stress signaling, critical gaps remain such as, the extent to which PFAS responses are tissue- and cell-type-specific, whether PFAS effects reflect general stress biology or compound-specific mechanisms, and the potential involvement of ribosomal stress and novel regulators in PFAS-mediated toxicity.

We hypothesize that PFOA and GenX elicit distinct, cell-type-specific responses in key stress pathways, including TGF-β-mediated fibrosis and p53-mediated DNA damage responses. By comparing responses across multiple tissues and pathways, we aim to uncover both shared and unique mechanisms of PFAS-induced toxicity, providing a mechanistic foundation for understanding tissue-specific health risks.

To better understand the effects of exposure to PFOA (perfluorooctanoic acid) and GenX in relation to DNA damage, cytotoxicity, and the potential for carcinogenic outcomes, two chemicals were evaluated across multiple cell lines. Key molecular pathways involved in stress responses were compared, including ribosomal stress, and TGF-β/SMAD signaling as outlined in Figure 1.

**Figure 1.**
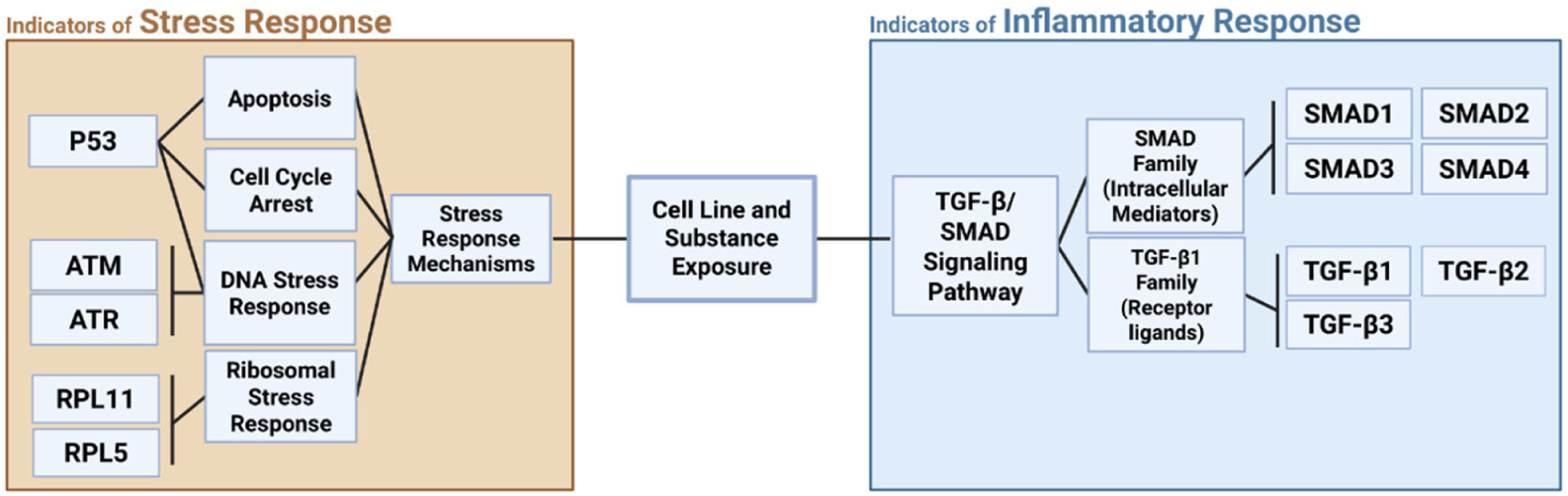
Key Genes and Pathways Linking DNA Damage Response, Ribosomal Stress, and TGF-β/SMAD Signaling in Tumor Microenvironment Regulation. This figure highlights key genes and proteins examined in the study. The DNA damage response is represented by p53, ATM, and ATR genes. RPL5 and RPL11, both ribosomal proteins, are involved in p53 regulation through the nucleolar stress response. The TGF-β/SMAD signaling pathway, a critical regulator of immune response and inflammation, is also depicted due to its role in modulating the tumor microenvironment.

## Methods

Detailed information about chemicals and materials used can be found in Appendices 1-3.

### Chemicals and test reagents

The ammonium salt form of PFOA, Pentadecafluorooctanoic acid ammonium salt (CAS No. 3825-26-1) of ≥98.0% purity was purchased from Sigma-Aldrich Corporation (St. Louis, MO, USA). The Undecafluoro-2-methyl-3-oxahexanoic acid form of GenX (CAS No. 13252–13-6) of 97% purity purchased from Synquest labs (Alachua, Florida, USA) was used in experiments.

Pentadecafluorooctanoic acid ammonium salt was dissolved in dimethyl sulfoxide (DMSO; CAS No. D8418) from Sigma-Aldrich to obtain a stock solution of 0.1 M. Largest equivalent volume of DMSO was spiked into the vehicle control experiment group. Different volumes of stock solution were added into the cell culture media to achieve the desired final concentrations of PFOA. It was empirically determined that 1 % v/v DMSO had negligible effect on cell response to Cytotoxicity was assessed using the Cell Counting Kit-8 (CCK-8, Dojindo Molecular Technologies, Cat. #CK04).

### Cell culture and analysis of cytotoxicity

Four adherent cell lines—A375 (melanoma), HepG2 (hepatocellular carcinoma), SN12C (renal cell carcinoma), and SW620 (colorectal cancer)—were used in this study. All cells were initially cultured in T25 flasks and subsequently transferred to T75 flasks at the third passage. The culture medium consisted of Dulbecco’s Modified Eagle Medium (DMEM) supplemented with 10% fetal bovine serum (FBS) and 1% penicillin-streptomycin (P/S). Flasks were maintained in a humidified incubator at 37 °C, 5% CO₂, pH 7.4, and 95% relative humidity. All media tested negative for mycoplasma contamination prior to experimentation. In addition, a sterility test was conducted in all experiments.

### Cytotoxicity Assay

Cytotoxicity was assessed by Cell Counting Kit-8 (CCK-8, Dojindo Molecular Technologies, Cat. #CK04) following the manufacturer’s protocol. Cells were seeded at a density of 4,000 cells per well in 96-well plates (Corning, Cat. #3599) and allowed to adhere for 24 hours in a humidified incubator (37°C, 5% CO₂). Stock solutions of perfluorooctanoic acid (PFOA, Sigma-Aldrich, Cat. #PFOA) and hexafluoropropylene oxide dimer acid (GenX, HFPO-DA, Sigma-Aldrich, Cat. #HFPO-DA) were prepared at concentrations of 0.1 M and 1 M, respectively, in dimethyl sulfoxide (DMSO, Sigma-Aldrich, Cat. #D8418), and stored at −20°C. Working solutions were freshly prepared by serial dilution in complete culture medium (Dulbecco’s Modified Eagle Medium with 10% fetal bovine serum and 1% penicillin-streptomycin; DMEM, Gibco, Cat. #11965-092) to achieve final concentrations of 0, 100, 250, 500, 750, and 1,000 μM for PFOA and GenX0, 500, 1,000, 2,000, 3,000, and 4,000 μM for GenX. The final DMSO concentration across all wells, including vehicle controls, was standardized to 1% (v/v) according to Wen et al^34^ and Yang et al.^35^

### Compound Treatment

Both PFOA and GenX were dissolved in DMSO at the final concentrations of 0.1M and 1M respectively and stored at -20°C.^36^ The control group was treated with complete medium containing 1% DMSO. Following a 48-hour incubation period at 37°C in a humidified 5% CO₂ atmosphere, 10 µL of the CCK-8 reagent was added to each well. Plates were incubated for a further 2 hours at 37°C, and absorbance was measured at 450 nm using a Varioskan™ LUX microplate reader (Thermo Fisher Scientific, Cat. #VLBLATD2). Wells containing blank culture medium with CCK-8 reagent (but no cells) were used to account for background absorbance.

### Quantification and Analysis

Cell viability was calculated as a percentage relative to vehicle-treated controls after subtracting the background absorbance. Dose-response cures were generated, and half-maximal inhibitory concentrations (IC_50_) were calculated using a four-parameter non-linear regression model in Graphpad Prism 9 (GraphPad Software, Version 9.5.0). Regression models were assessed for goodness-of-fit (R^2^ values). Experiments were performed as the mean ± standard deviation (SD).

### GenX Immunoblotting

Immunoblotting was carried out as described previously (Hu et al., 2020). To standardize protein concentrations among lysates, the Enhanced BCA Protein Assay Kit was used and a total of 20 - 30 uL of total protein were loaded in SDS-PAGE gels (Appendix 2), and electrophoresis of samples was performed in a gel electrophoresis system with running buffer (Appendix 2) along a protein size marker (10-180 kDa). Separated proteins were transferred to a PVDF membrane using a wet transfer system and transfer buffer (Appendix 1). Resulting membranes were blocked on an orbital shaker for 1h at room temperature by adding a blocking buffer (5% BSA dissolved in TBST, Appendix 2). and incubated with primary and secondary antibodies diluted in TBST containing 1% BSA, with ratios in Appendix 3. The membranes were imaged using a chemiluminescence imaging system.

### RNA extraction and cDNA synthesis

Total RNA extraction using Trizol reagent has been described in detail in previous research and follows standard protocols.^37^ For cDNA synthesis, the HiScript II 1st Strand cDNA Synthesis Kit was used. RNA concentration was estimated using Nanodrop. First-strand cDNA synthesis reaction solutions were assembled in RNase-free centrifuge tubes with a total of 1 μg RNA per sample. The reaction solutions were mixed gently and sequentially reacted at 25°C for 5 min, 50°C for 15 min, and 85°C for 2 min in a PCR thermal cycler.

### Quantitative PCR

All quantitative (q)PCR experiments were performed using 96-well PCR plates. In the 20 μL reaction system, 10 μL was ChamQ Universal SYBR qPCR Master Mix, 0.4 μL of 1 μM forward primer, 0.4 μL of 1 μM reverse primer, 7.2 μL of RNase-free ddH_2_O, and 2 μL of template cDNA. Primer details are shown in Appendix 4. The qPCR thermal cycler was set as 95℃ for 30 seconds with 1 repetition (Pre-denaturation), 95℃ 10 seconds and 60℃ 30 seconds with 40 repetitions (Cycling Reactions), and a melting curve acquisition (Default melting curve program). Relative expression was calculated using the 2^-ΔΔCt method.

### Statistical analysis

All cytotoxicity results are presented as a value and 95% confidence interval while qPCR results are presented as mean ± standard deviation (SD). Data were analyzed by two-tailed t-tests for groups with 2 variables and analysis of variance (ANOVA) for groups with more than 2 variables. GraphPad Prism version 9.5.1 for Windows was used. P-values < 0.05 were considered significant.

## Results

### Cytotoxicity assessment

All cells were more tolerant to GenX and displayed similar trends in cytotoxicity assessment shown below in Figure 2. IC_50_’s ranged from 213.2 μM to 5048 μM, with the highest sensitivity observed in A375 and SW620 cells and lower sensitivity in HepG2 and SN12C cells. The A375 cell line showed a high sensitivity to PFOA with an IC_50_ value of 213.2 μM (R^2^ = 0.9375) (Fig. 1A) and more tolerance to GenX with an IC_50_ value of up to 1555 μM (R^2^= 0.9920) (Fig. 1B). HepG2 cells were moderately low in sensitivity to PFOA with an IC_50_ of 527.9 μM (R^2^ = 0.9974) (Fig. 1A). Overall, all cell types were more resistant to GenX exposure with an average estimated IC_50_ value of 2377 μM (R^2^ = 0.9747) (Fig. 1B). SN12C cells showed the most resistance overall, with an IC_50_ value of 616.3 μM for PFOA and IC_50_ values up to 5,048 μM (R^2^ = 0.9400) for GenX (Fig. 1B). The SW620 cell line exhibited moderate sensitivity to PFOA with an IC_50_ of 271.2 μM (R^2^ = 0.9965) (Fig. 1A). Similarly, SW620 also showed moderate tolerance to GenX with an IC_50_ of 2126 μM (R^2^ = 0.9874) (Fig. 1B).

**Figure 2:**
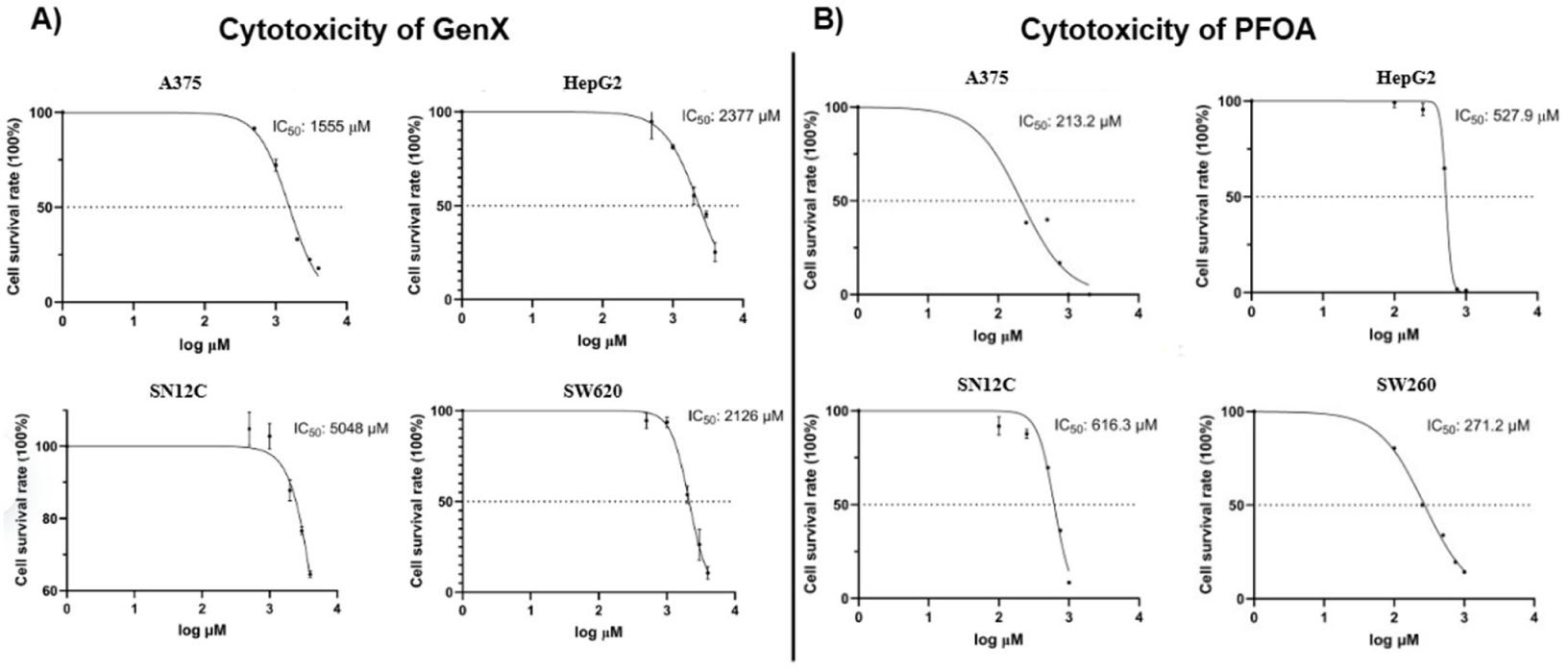
Viability of A375, SN12C, SW620, and HepG2 cells. A) Cytotoxicity to PFOA and B) Cytotoxicity to GenX. The data obtained through CCK-8 experiments were transformed to a log base after processing, fitted using nonlinear regression, and resulted in cell viability curves for each cell in response to PFOA concentration. Dashed lines represent the PFOA concentration at which half of the cells survived (IC_50_), and the best-fit values are labeled in the upper right corner.

As displayed in Table 1, IC_50_ values were obtained by CCK-8 assay and estimated by fitting the results using nonlinear regression with a 95% confidence interval range.

**Table 1:** IC_50_ of A375, HepG2, SN12C, and SW620 cell stress response to PFOA and GenX.

| Cell lines | PFOA treated $IC_{50}$ μM (95% CI) | GenX treated $IC_{50}$ μM (95% CI) |
| --- | --- | --- |
| A375 | 213.2 [23.54, 393.8] | 1555 [1452, 1664] |
| HepG2 | 527.9 [514.6*, 541.2] | 2377 [2148, 2630] |
| SN12C | 616.3 [558.8, 670.1] | 5048 [4451, 6193**] |
| SW620 | 271.2 [239.8, 304.4] | 2126 [1984, 2267] |

\* Unable to estimate specific values
\*\* Unable to estimate specific values due to exceeding limit set

### Cell Line Exposure effects on Inflammatory and Stress Responses Focusing on TGF-β/SMAD Pathway

Having established acute cytotoxicity estimates for PFOA and GenX, we sought to understand the molecular consequences of exposure, with focus on DNA damage/repair (ATM/ATR, P53 expression), ribosomal stress (RPL5/11) and immune signaling (TGF-β/SMAD pathway). We found effects to differ between the two chemicals as well as between cell lines (Figs. 3, 4, 5, 6). Changes in gene expression and protein level are summarized in Table 2.

**Figure 3.**
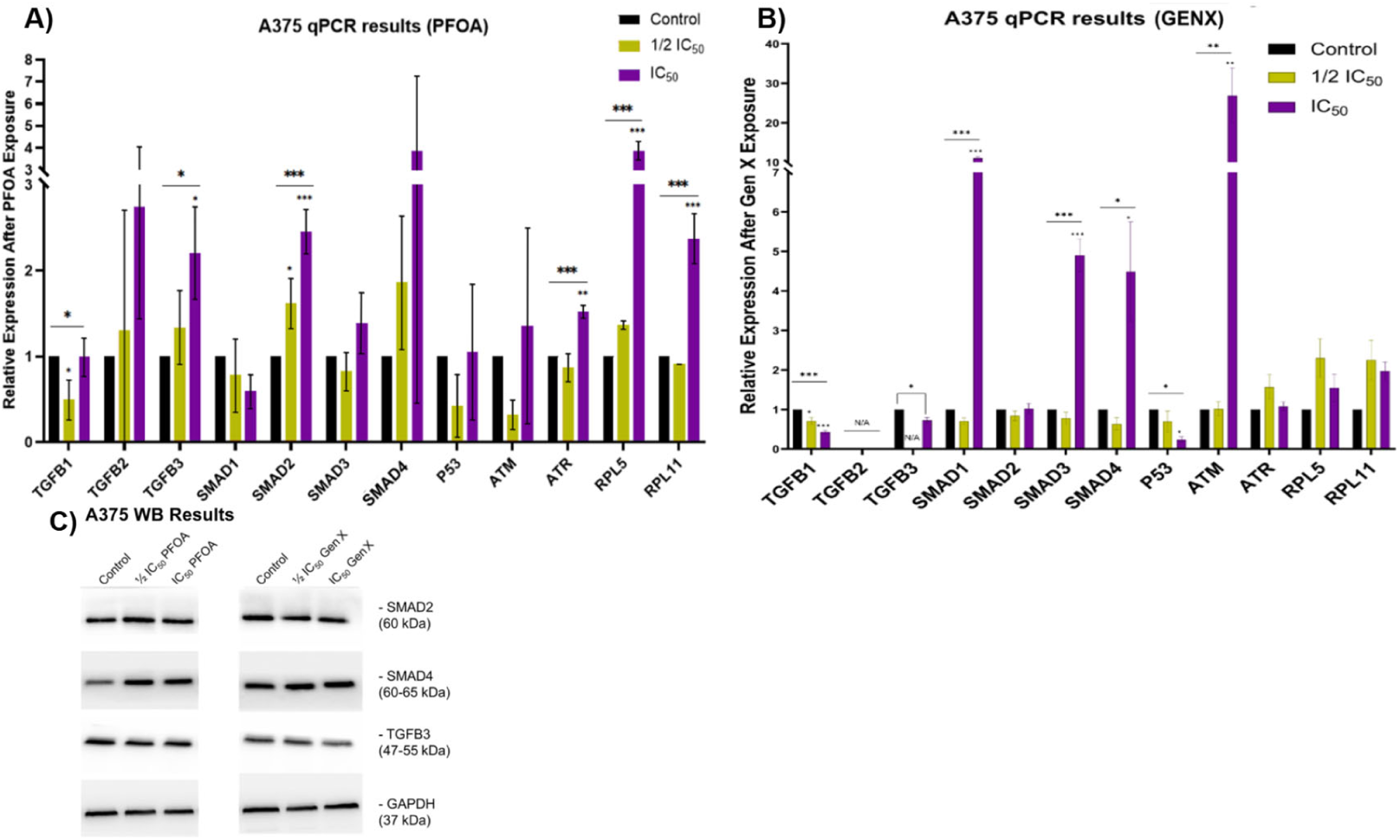
Altered gene and protein expression in A375 cells following PFOA or GenX treatments. A375 cells were treated with vehicle, A) PFOA or B) GenX with concentration of 1/2 IC_50_, IC_50_ for 48 h, and relative mRNA levels were determined by RT-qPCR. *, p < 0.05; **, p < 0.01,*** p < 0.001 C) The WB results of target inflammatory response proteins extracted from the control group and PFOA or GenX treated A375 cells with concentration of 1/2 IC_50_, IC_50_.

**Figure 4.**
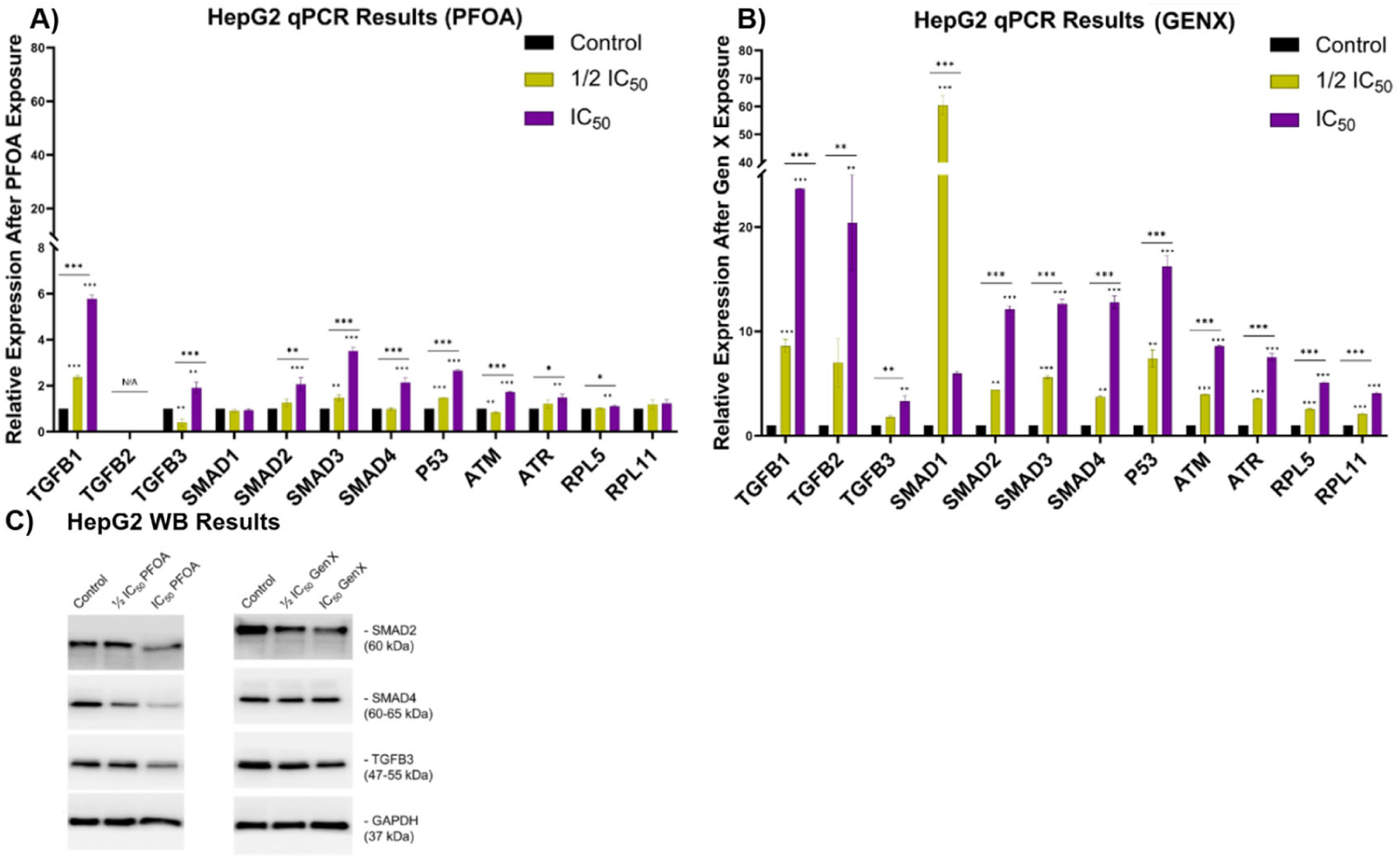
Altered gene and protein expression in HepG2 cells following PFOA or GenX treatments. HepG2 cells were either mock-treated or treated with A) PFOA or B) GenX with concentration of 1/2 IC_50_, IC_50_ for 48 h and relative mRNA levels were determined by q-PCR. *, p < 0.05; **, p < 0.01, *** p < 0.001; C) The WB results of target inflammatory response proteins extracted from the control group and PFOA or GenX treated HepG2 cells with concentration of 1/2 IC_50_, IC_50_.

**Figure 5.**
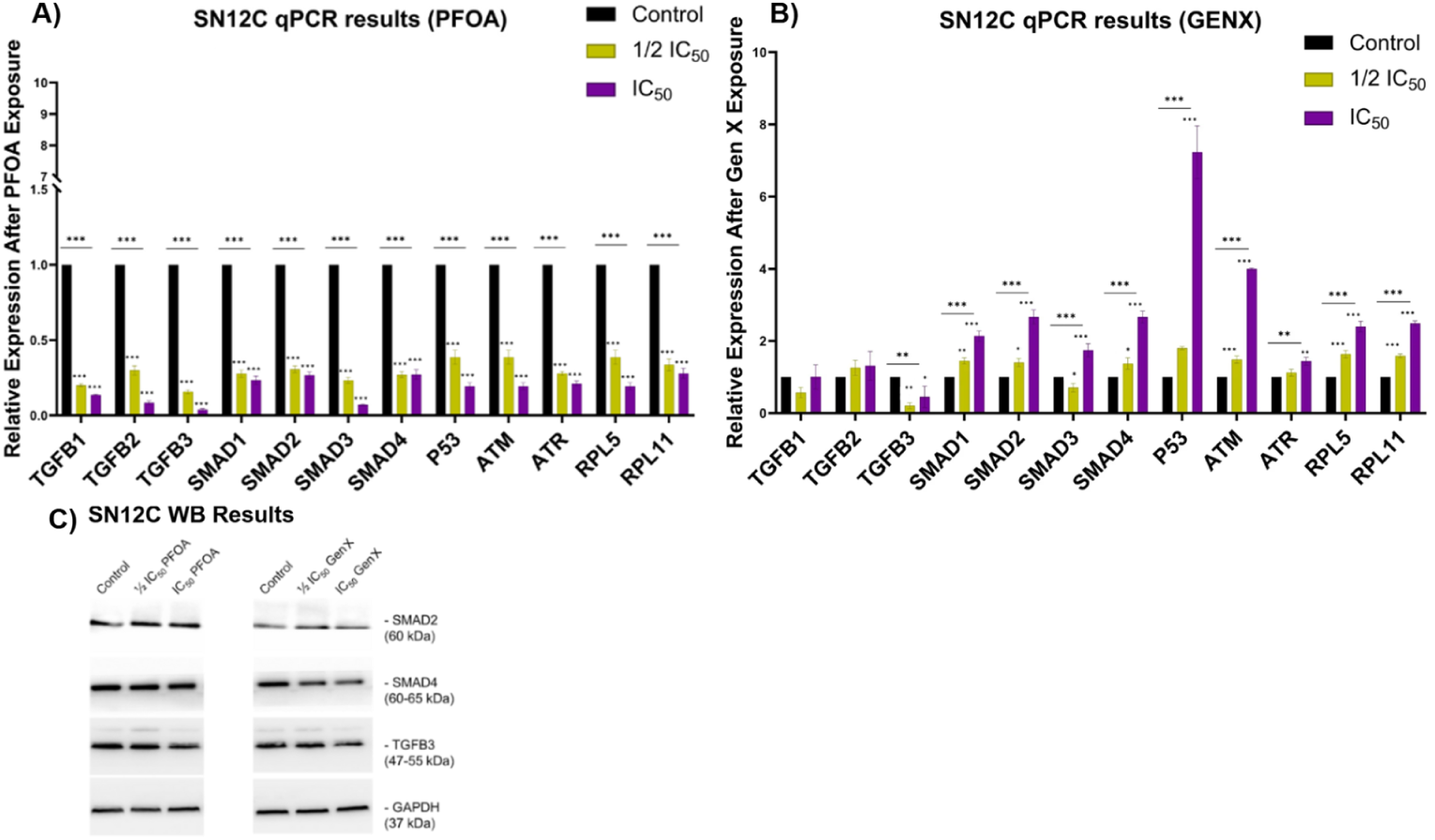
Altered gene and protein expression in SN12C cells following PFOA or GenX treatments. SN12C cells were either mock-treated or exposed to A) PFOA or B) GenX at concentrations of ½ IC_50_, IC_50_ for 48 h. Relative mRNA levels were quantified by q-PCR. \*, p < 0.05; \*\*, p < 0.01; \*\*\* p < 0.001. (C) Western Blot analysis of target inflammatory response proteins extracted from control (mock-treated) and PFOA- or GenX-treated SN12C cells with concentration of ½ IC_50_, IC_50_.

**Figure 6.**
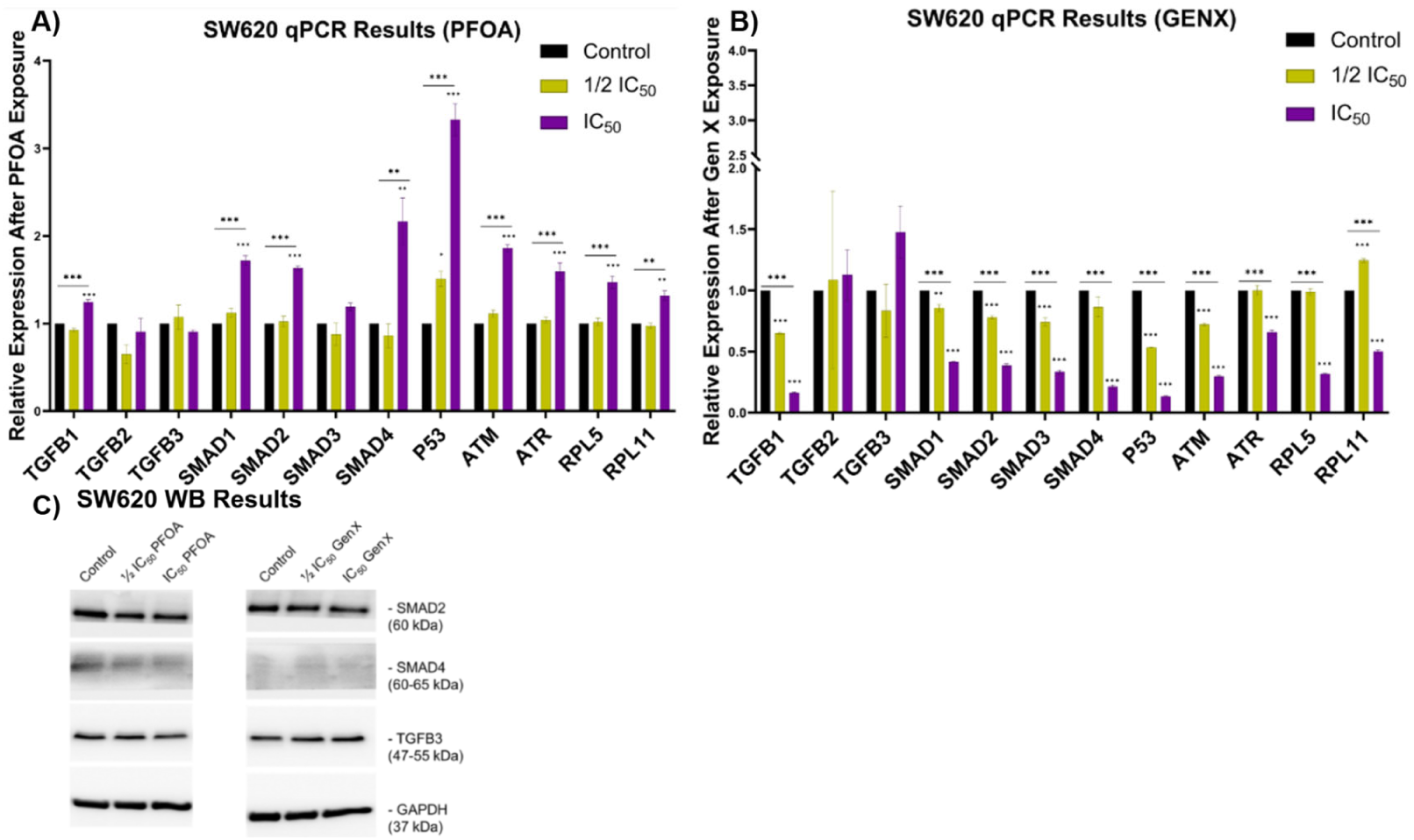
Altered gene and protein expression in SW620 cells following PFOA or GenX treatments. SW620 cells were either mock-treated or treated with A) PFOA or B) GenX with concentration of 1/2 IC_50_, IC_50_ for 48 h and relative mRNA levels were determined by q-PCR. *, p < 0.05; **, p < 0.01, *** p < 0.001; C) The WB results of target inflammatory response proteins extracted from the control group and PFOA or GenX treated SW620 cells with concentration of 1/2 IC_50_, IC_50_.

**Table 2:**
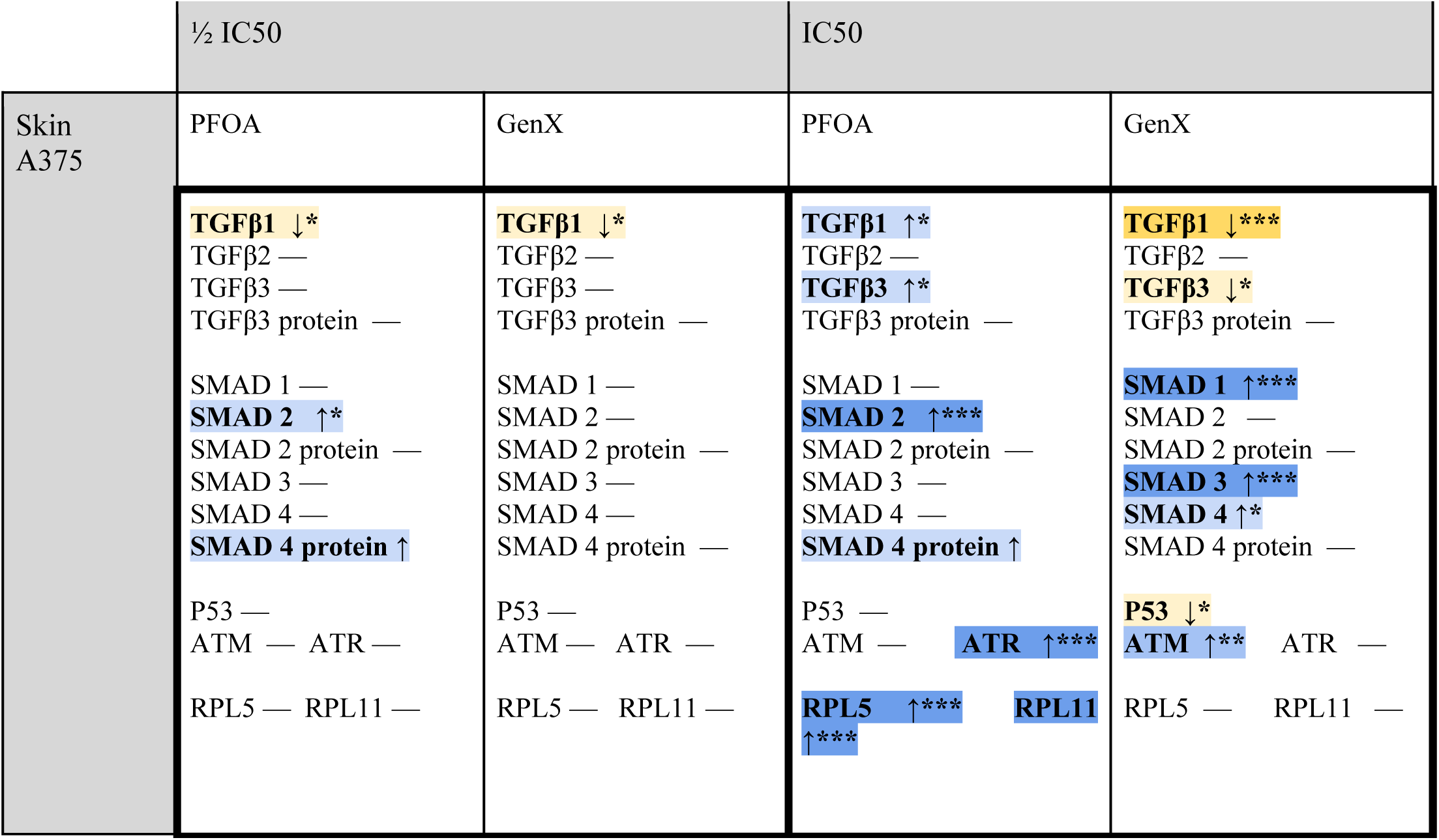
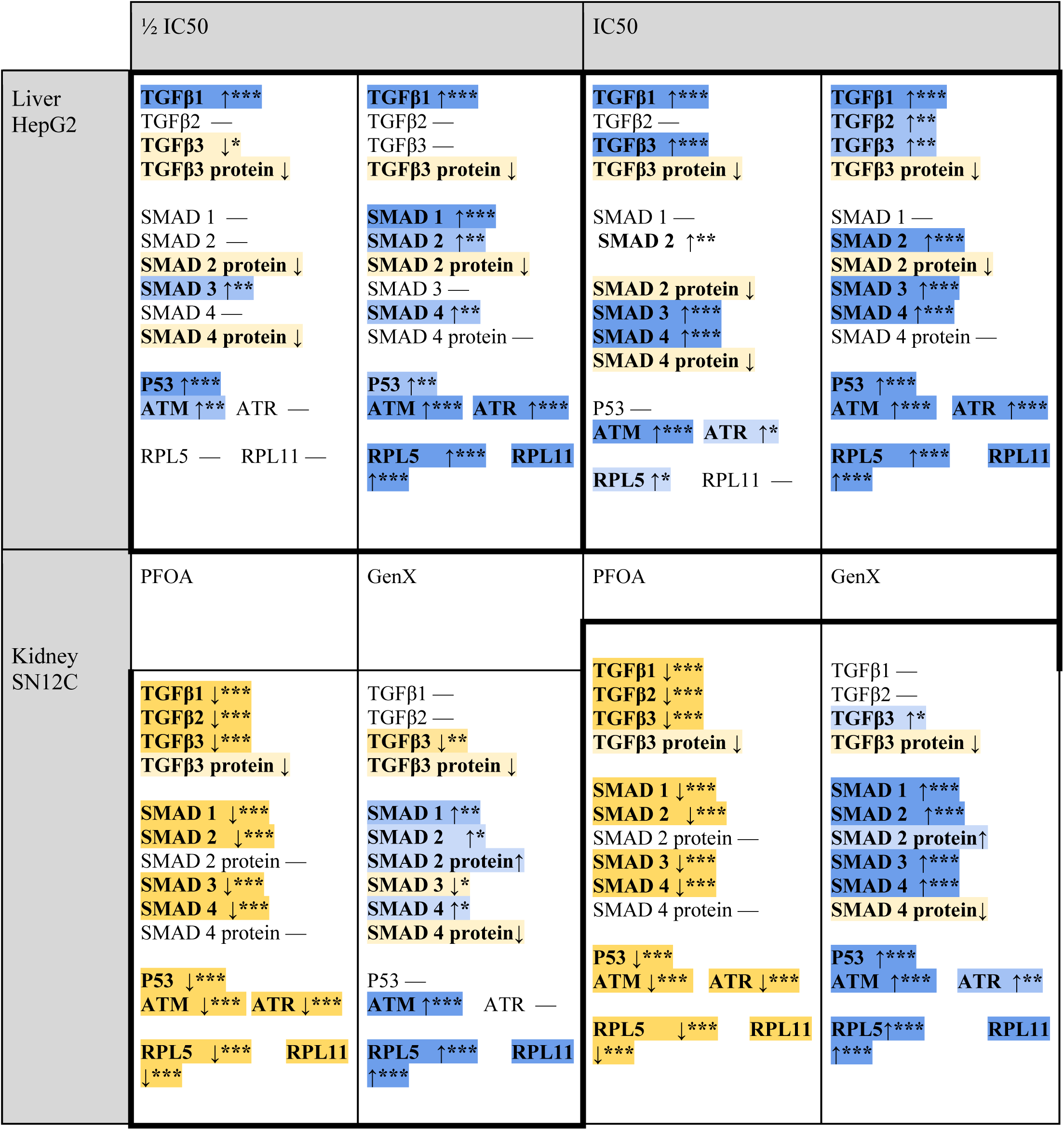
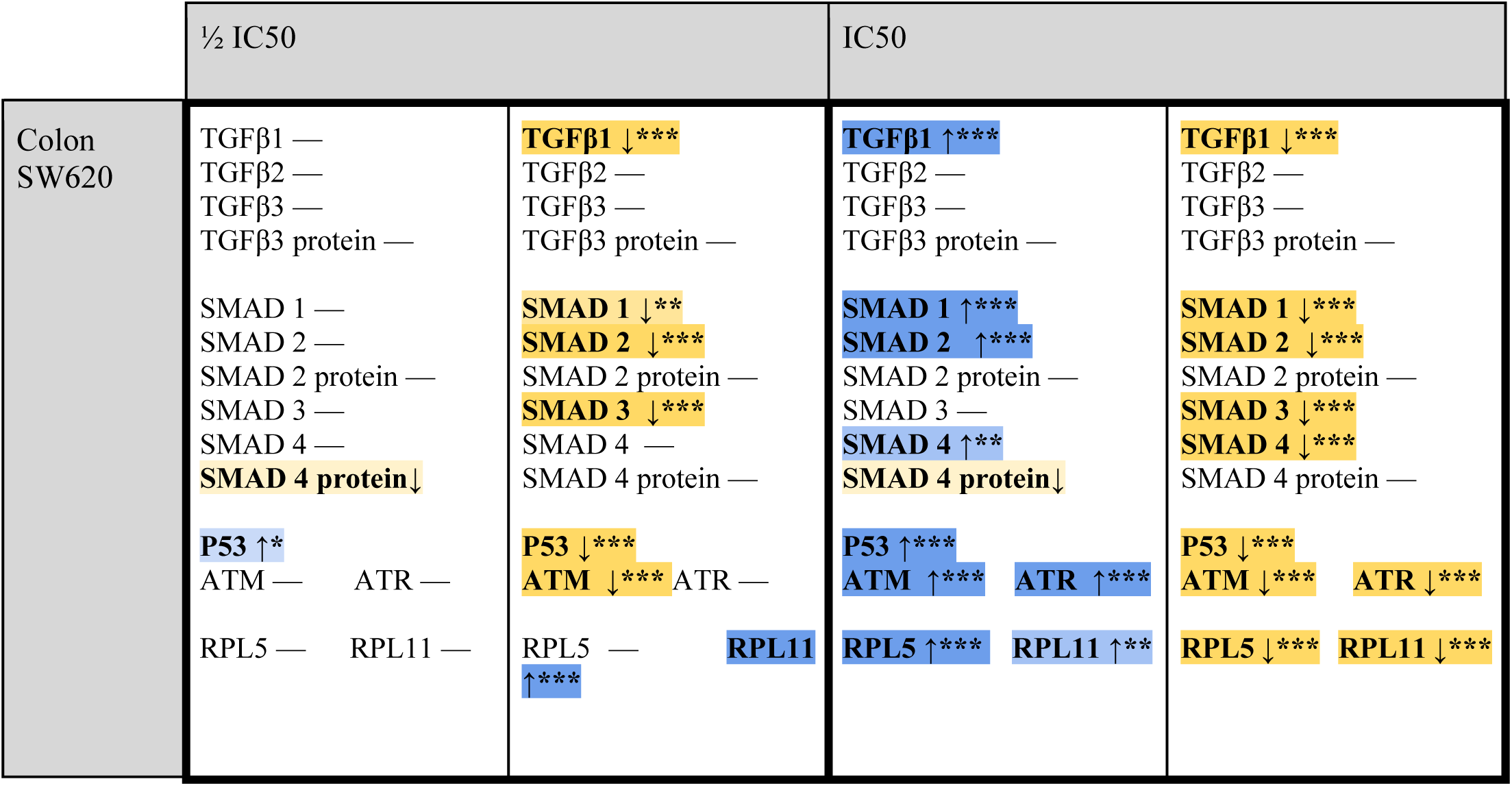
Summary of Gene and Protein Expression of Inflammatory and DNA Stress Response Markers. This table summarizes the changes in gene expression (measured by qPCR) and protein levels (measured by immunoblot/western blot) of target inflammatory and stress response markers. Unless otherwise specified with “protein”, entries in the table refer to the gene expression. *, p< 0.05; **, p < 0.01, *** p < 0.001. Yellow indicates a decrease while blue indicates an increase in expression.

#### i) A375 cell line

Exposure of A375 cells to PFOA increased SMAD2 gene expression with p <0.05 (Fig. 3A) and SMAD4 protein level in both ½ IC_50_ and IC_50_ concentrations, while no significant trend was observed for the expression of other proteins (Fig. 3C). As shown in Figure 3A, more significant changes in the expression of genes occurred at the IC_50_. At ½ IC_50_, TGFβ_1_ gene expression decreased (p < 0.05), while SMAD2 gene expression increased. At IC_50_, TGFβ_1_ gene increased on the contrary (p < 0.05), along with increases in TGFβ_3_ (p < 0.05), SMAD2, ATR, RPL5, and RPL11 (p < 0.001) compared to untreated controls. However, the expression of other genes such as p53, ATM, TGFβ_2_, TGFβ_3_, SMAD1 and SMAD3 did not change significantly in the experimental group. In the GenX treated A375 cells, the expression of TGFβ_1_ was decreased at ½ IC_50_ concentration compared to the control (Fig. 3B). At ½ IC_50_ there were no other significant gene expression or protein level changes. At IC_50,_ there were more changes again. In the TGF-β family genes, TGFβ_1_ gene expression decreased at IC_50_-treated concentrations (p<0.001), and the expression of TGFβ_3_ increased significantly at both ½ IC_50_ and IC_50_ concentrations (p < 0.05 and p < 0.001), while TGFβ_2_ did not show a corresponding signal in the experiment (N/A). In the SMAD family genes, SMAD1, SMAD3 and SMAD4 showed significant up-regulation (p < 0.001 for SMAD1, SMAD3 and p < 0.05 for SMAD4), which is consistent with the activation of the TGF-β signaling pathway. The expression of SMAD2 did not show statistically significant changes after GenX treatment. In addition, ATM also showed a significant increase in expression at IC_50_ concentrations (p < 0.01), suggesting that a DNA damage response mechanism may have been induced in the presence of GenX. However, the p53 gene showed a decrease in expression at IC_50_ concentration (p < 0.05).

#### ii) HepG2 cell line

In the HepG2 liver cell line of Figure 4, there were significant changes caused by both chemicals at 1/2 IC_50_ and IC_50_ levels. These effects were generally similar between PFOA and GenX.

For exposure to PFOA at 1/2 IC_50_ the expression of the TGFβ_1_ gene was more than 2-fold higher (p<0.001). SMAD3 (p<0.01), p53 (p<0.001), and ATM (p<0.01) gene expression also increased. Both TGFβ_3_ gene (p<0.05) and proteins expressed less at ½ IC_50_ as well as SMAD2 and SMAD4 protein levels (Fig. 4C). At IC_50_ the TGFβ_1_ gene expression increased again to around 6-fold higher than the control (p<0.001). SMAD2 (p<0.01), SMAD3 (p<0.001), SMAD4 (p<0.001), ATM (p<0.001), ATR (p<0.05), and RPL5 (p<0.05) gene expressions also increased. Interestingly, TGFβ_3_, downregulated at ½ IC50, increased to 2-fold compared to the control group at IC50 (p < 0.001). However, the TGFβ_3_ protein levels decreased, along with SMAD2 and SMAD4 proteins, despite their respective gene upregulations. The expression data for TGFβ_2_ were not applicable to this analysis as the signal from the control could not be observed (N/A).

In the GenX treatment group (Fig. 4B), all target gene expressions showed increases in different degrees. Specifically, both TGFβ_1_ and TGFβ_2_ expressed more than 20-fold higher at IC_50_ (p<0.001 and p<0.01 respectively). The SMAD1 expression showed the greatest increase at the ½ IC_50_ concentration (p<0.001), but its increase at the IC_50_ concentration is not statistically significant. At ½ IC_50_ TGFβ_1_ (p<0.001), SMAD1 (p<0.001), SMAD2 (p<0.01), SMAD4 (p<0.01), p53 (p<0.01). ATM (p<0.001), ATR (p<0.001), RPL5 (p<0.001), RPL11 (p<0.001) gene expression increased, while the TGFβ3 and SMAD2 proteins decreased. At IC_50_ TGFβ_1_ (p<0.001), TGFβ_2_ (p<0.01), TGFβ_3_ (p<0.01), SMAD2, SMAD3, SMAD4, p53, ATM, ATR, RPL5, and RPL11 (all p<0.001) gene expression increased. TGFβ3 and SMAD2 proteins showed a decreasing trend with increasing GenX concentration.

In the SN12C cell line of Figure 5, the expression of genes associated with TGF-β pathway, cellular stress response, DNA damage repair, ribosome assembly, and immune signaling all significantly decreased after PFOA treatment with both the ½ IC_50_ and IC_50_ concentrations of PFOA, and the expression of these genes was particularly more pronounced at IC_50_ concentrations, with the statistical significance reaching the level of p < 0.001 (Fig. 5A). Notably, despite decreases in gene expression, SMAD2 and SMAD4 protein concentrations didn’t change. In contrast, under GenX treatment (Fig. 5B), all genes other than TGFβ_1_, TGFβ_2_,and TGFβ_3_ all showed a significant increase in expression at IC_50_ concentration (p < 0.001). SMADs, ATM/ATR, RPL5/RPL11, and especially p53 showed a dose-dependent increase trend with increasing GenX. However, with the TGF-β family, TGFβ_3_ was decreased in expression at ½ IC_50_ and IC_50_ concentrations from the control, while TGFβ_1_ and TGFβ_2_ were unchanged. There was also an upward trend in SMAD2 protein levels and a decreased trend in SMAD4 protein levels in response to escalating GenX concentrations.

#### iii) SN12C cell line

In the SW620 cell line of Figure 6, p53 mRNA was significantly upregulated, and SMAD4 protein levels decreased. At IC_50_ SMAD4 protein levels continued to decrease, but the TGFβ_1_ (p < 0.001), SMAD1 (p < 0.001), SMAD2 (p < 0.001), and SMAD4 (p < 0.01) genes exhibited increased expression at the IC_50_ concentration. Levels of p53, ATM, ATR, RPL5, and RPL11 mRNAs were increased by treatment with PFOA at the IC_50_ concentration.

#### iv) SW620 cell line

In the GenX treatment group (Fig. 6B), the target gene expressions showed an overall decreasing trend. Within the TGF-β family, there was a significant decrease in TGFβ_1_ in both groups (p<0.001). The expression of TGFβ_2_ and TGFβ_3_ increased but did not show statistical significance. The expression of SMAD family genes was also significantly affected by GenX treatment. At ½ IC_50_ a decrease in TGFβ_1,_ SMAD1, SMAD2, SMAD3, p53, and ATM, while RPL11 increased (p<0.001 for all except p<0.01 for SMAD1). At IC_50_, maybe gene expressions continued to decrease, such as TGFβ_1_, SMAD1, SMAD2, SMAD3, SMAD4, p53, ATM, ATR, and RPL5 (p<0.001 from control for all). After increasing with exposure to ½ IC_50_, RPL11 decreased to be below the levels of the control with IC_50_ exposure (p<0.001).

## Discussion

### PFOA and GenX cytotoxicity

GenX, developed as a safer alternative to PFOA,^18^ exhibited higher IC₅₀ values across all tested cell lines, indicating lower cytotoxicity. This observation is consistent with prior studies showing greater cytotoxicity for PFOA in immune cells such as THP-1 monocytes and Jurkat T-cells^38^. Notably, contrasting results have been reported in thyroid models, where GenX demonstrated greater toxicity than PFOA,^39^ pointing to tissue- and lineage-specific variability in PFAS response. GenX exhibits substantially higher IC₅₀ values (1555–5048 µM) compared to PFOA (213–616 µM), indicating lower cytotoxic potency under the conditions tested. Rat studies confirm PFOA’s higher potency at matched external doses yet suggest that internal target-organ dosimetry can equalize the effects of PFOA and GenX.^16^ Therefore, our current findings should be contextualized as part of a broader toxicological framework that integrates both cytotoxicity and molecular response.

In this study, we selected HepG2, SN12C, SW620, and A375 cell lines based on known PFAS tissue distribution and toxicokinetic considerations. The liver is a primary target for PFAS accumulation, while kidneys play a key role in excretion, the colon reflects exposure through enterohepatic circulation, and skin represents systemic distribution and potential local effects. These choices are supported by prior studies reporting measurable PFAS concentrations in these tissues in both animal and human models.^11,40,41^ While our study focused on tissues known to accumulate PFOA, future work including immune-based cells will be important to determine whether these are more directly susceptible to PFAS-induced immunotoxicity, as suggested by prior reports of PFAS-associated immune suppression.^42,43^

We also note that all four cell lines used are cancer-derived and immortalized, which may influence metabolic capacity, stress responses, and susceptibility to PFAS exposure. While these models provide valuable mechanistic insight, their responses may not fully reflect those of normal, non-cancerous tissues in vivo. Thus, the generalizability of our findings to non-immortalized cells within an organism is limited. Future studies using primary human cells or organotypic models will be important to validate whether the observed mechanisms operate in physiologically relevant tissues across different organ systems.

### DNA Damage and p53 Pathway Activation

A key aim of our study was to examine whether the observed cytotoxicity was underpinned by genotoxic stress and DNA damage. Although DNA damage markers appeared more dynamically regulated in metabolically active cells, this observation likely reflects differences in cellular stress response capacity rather than metabolic transformation of PFOA, as PFAS are generally considered resistant to biotransformation with only limited metabolism reported for certain precursors.^44^ This pattern was echoed in the expression of upstream DNA damage regulators. In HepG2 cells, both PFOA and GenX upregulated ATM, ATR, and p53—central components of the DNA damage sensing and repair machinery. However, tissue-specific divergence emerged: in SN12C cells, PFOA suppressed ATM/ATR and p53 expression, while GenX activated them. In the colon, the inverse pattern was observed, with PFOA inducing ATM/ATR/p53 upregulation and GenX causing downregulation. These discordant findings suggest chemical-specific differences in how DNA damage is detected and processed across tissues and warrant further studies exploring post-translational modifications of p53,^45^ including phosphorylation and acetylation, to better clarify its activation status.

### Ribosomal Stress as a p53 Modulator

Given the interplay between ribosomal dysfunction and p53 activation, we also evaluated expression of ribosomal stress sensors RPL5 and RPL11. In A375 cells, both genes were significantly upregulated only under IC₅₀ PFOA treatment, suggesting a stress threshold that triggers the nucleolar surveillance pathway. In contrast, HepG2 cells exhibited a clear and consistent increase in RPL5 and RPL11 across both treatments, corresponding with p53 upregulation and reinforcing the role of ribosomal stress in hepatic PFAS response.

SN12C and SW620 cell lines again showed inverse responses to the two compounds. In SN12C cells, PFOA suppressed RPL5/RPL11 expression, while GenX upregulated both. In SW620 cells, PFOA elevated ribosomal stress markers, while GenX reduced them. These observations suggest that while GenX may more strongly disrupt ribosome homeostasis in the kidney-derived cell line, PFOA exerted greater ribosomal stress in the colon-derived cell line, possibly through differential impacts on nucleolar integrity or translation control.

### TGF-β/SMAD Signaling and Inflammation

This study provides the first comparative analysis of TGF-β/SMAD pathway modulation by the perfluoroalkyl substances PFOA and GenX across HepG2, SN12C, and SW620 cell lines. TGF-β signaling is a pivotal regulator of inflammation, fibrosis, and epithelial-to-mesenchymal transition (EMT), and its dysregulation has been implicated in both endocrine disruption and immune dysfunction.^46,47,48^ In HepG2 cells, both compounds induced robust transcriptional upregulation of SMAD2 and SMAD4; however, corresponding protein levels remained unchanged, suggesting that PFAS exposure may perturb translational efficiency or promote post-translational degradation, potentially via proteasomal or miRNA-mediated mechanisms.

Notably, distinct responses were observed in SN12C and SW620 models. In SN12C cells, PFOA suppressed TGF-β/SMAD gene expression, while GenX led to significant induction, suggesting possible compound-specific effects on renal inflammation or repair pathways. Conversely, in SW620 cells, PFOA enhanced SMAD expression, whereas GenX inhibited it—indicating opposing regulatory dynamics that may reflect distinct cellular stress adaptations. These variations underscore the tissue-specific complexity of PFAS-induced signaling responses.

Canonical TGF-β signaling is initiated by ligand binding (TGF-β1, TGF-β2, or TGF-β3), leading to phosphorylation of receptor-regulated SMADs (R-SMADs, including SMAD2, SMAD3, and in some contexts, SMAD1), which then associate with the common-mediator SMAD4 and translocate to the nucleus to regulate target gene expression. In our study, transcriptional induction of SMADs coincided with marked upregulation of TGF-β1 and increased expression of DNA damage response markers such as p53, ATM, and ATR. This co-activation suggests that SMAD signaling may be engaged in response to PFAS-induced genotoxic stress, coordinating cell cycle arrest with pro-apoptotic or DNA repair programs.Intriguingly, despite the elevated transcript levels, SMAD2 and SMAD4 protein expression declined at higher toxin concentrations, consistent with possible post-transcriptional repression or proteolytic turnover under sustained stress. These findings align with the established profibrotic and oncogenic roles of TGF-β/SMAD signaling in chronic hepatic injury and support growing concerns regarding the hepatotoxic potential of both legacy and emerging PFAS.

It is important to note that the concentrations applied in this study represent nominal, administered doses, which may differ from the actual free concentrations present in culture media or internalized by cells. PFAS are known to bind serum proteins, which can lower bioavailable fractions. While direct quantification of media or intracellular concentrations was not conducted here, previous studies have shown that such factors can significantly affect effective dosimetry.^49^ Future work measuring post-exposure media concentrations (e.g., LC–MS/MS) or applying physiologically based pharmacokinetic (PBPK) modeling will be critical to better align in vitro exposures with internal target-organ dosimetry.

Our study design, however, provides an internal comparative framework that helps to mitigate this limitation. By exposing multiple cell types (endothelial and endometrial) to the same nominal concentrations, we were able to identify differential responses across models. In this context, each cell line effectively serves as a comparator or control for the others, highlighting relative susceptibilities and pathway-specific effects even in the absence of direct dosimetry measurements.

To clarify upstream regulation of DNA damage signaling, we specifically note that ATM and ATR kinases, together with p53, act as central regulators of the DNA damage response. These proteins govern recognition and repair of DNA double-strand breaks.

## Conclusion

Taken together, our results demonstrate that while GenX exhibits lower overall cytotoxic potency than PFOA under the in vitro conditions tested, it nonetheless induces distinct effects on DNA integrity, ribosomal stress, and inflammatory signaling that are both tissue- and pathway-specific. These findings highlight that reduced cytotoxicity does not equate to biological inertness and challenge assumptions regarding universal safety of PFAS replacement compounds. Importantly, the observed differences in cellular responses likely reflect intrinsic toxicodynamic properties rather than accumulation effects, which cannot be directly assessed within the present in vitro model.

Beyond cellular context, PFAS exposure remains a significant public health concern. Environmental and epidemiological studies have reported PFAS blood concentrations exceeding established Tolerable Daily Intakes (TDIs), particularly in populations exposed through contaminated drinking water and agricultural systems.^22^ In this broader environmental setting, physiochemical properties such as polarity and functional group substitution may influence environmental persistence and distribution-for example, GenX exhibiting greater mobility in aqueous systems, while PFOA shows stronger association with lipophilic tissues and soils.^5,7^ Future studies should integrate systems-level approaches—combining transcriptomic, proteomic, and phospho-proteomic analyses—to better assess the interplay between DNA damage responses, ribosomal integrity, and inflammatory signaling across PFAS classes. In particular, post-translational modifications of p53 and SMAD family members may provide crucial insight into PFAS-specific toxicodynamic pathways and support more informed risk assessment, regulatory decisions, and development of safer chemicals.

**Appendix 1:**
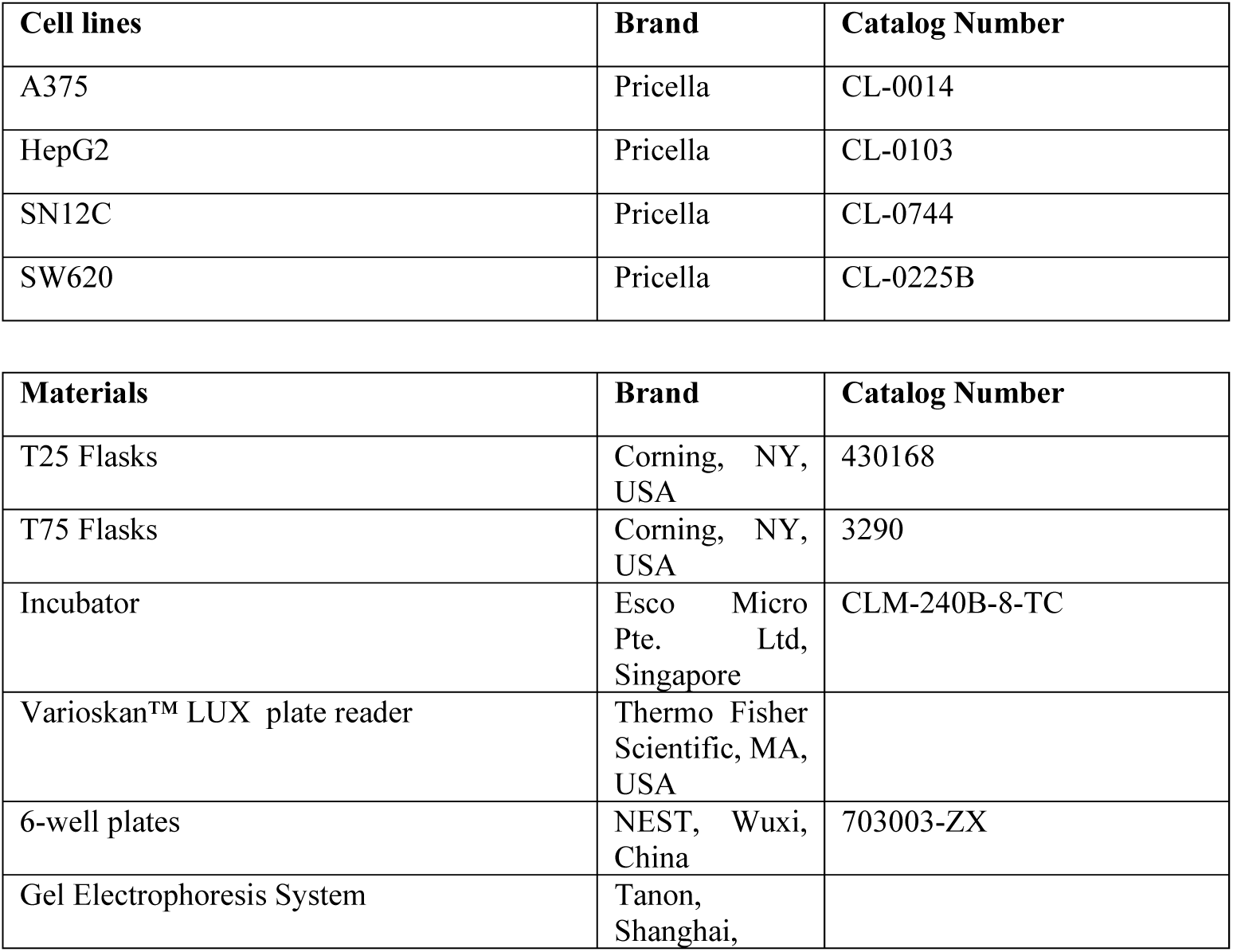
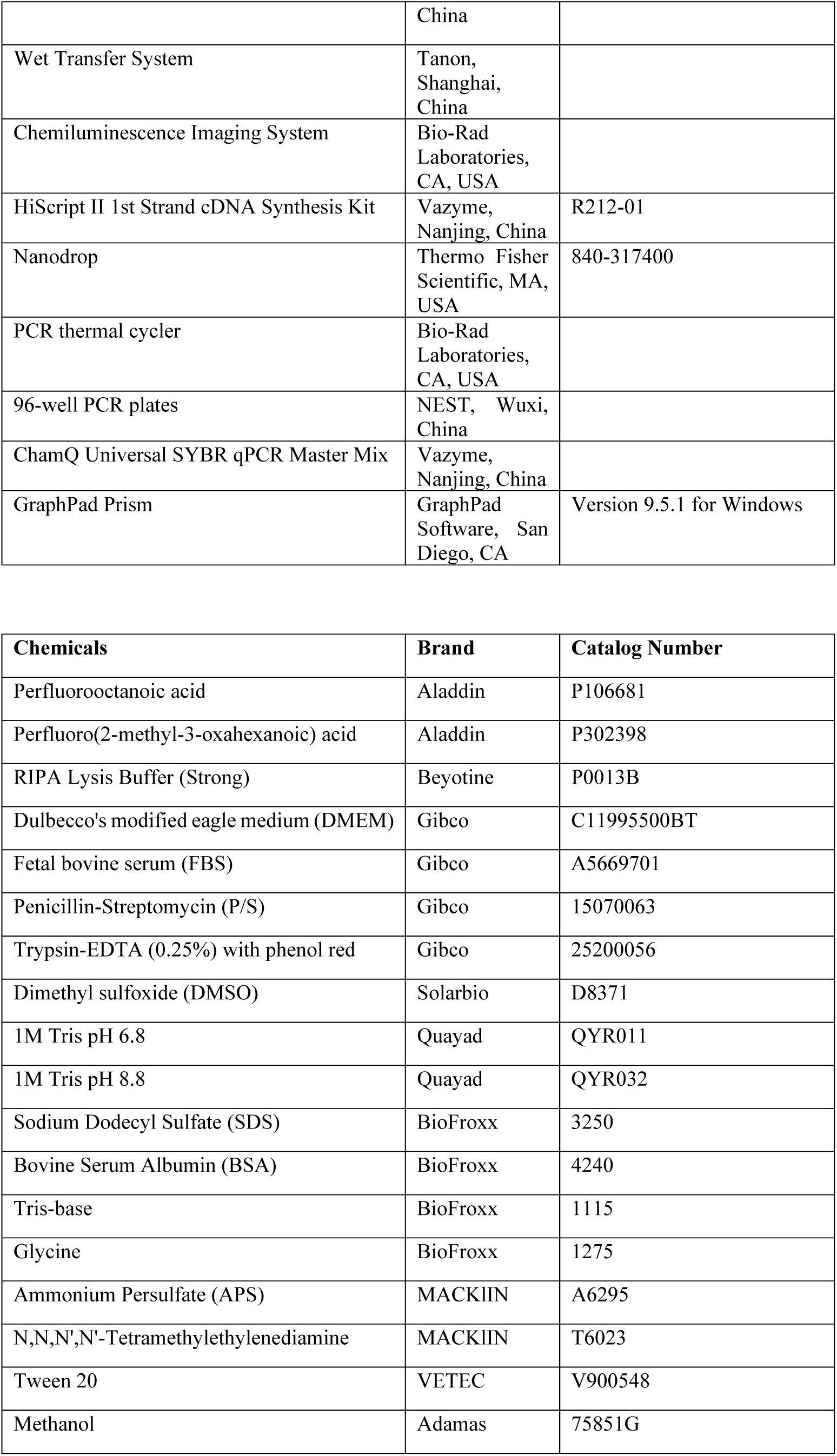
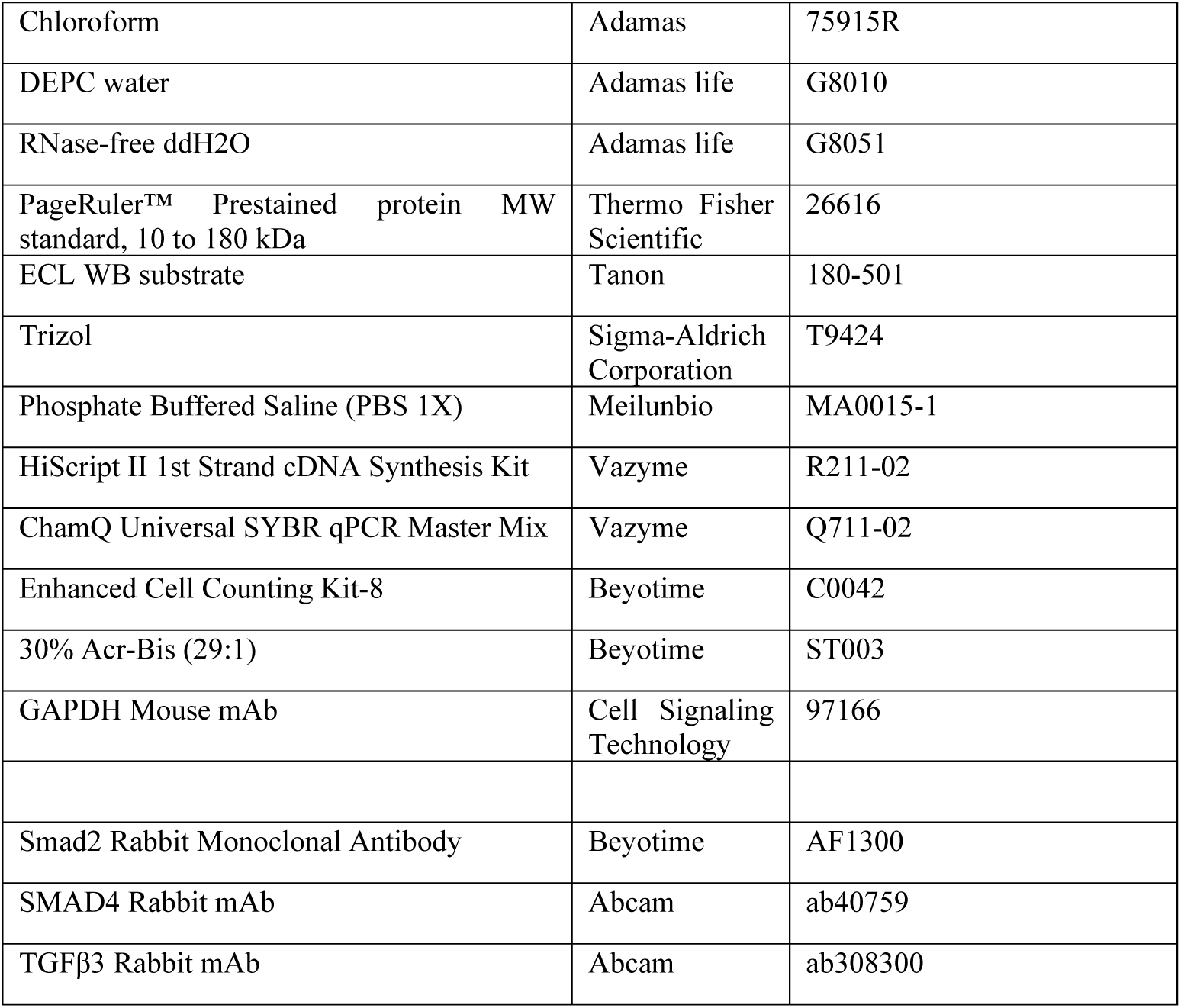
List of materials.

**Appendix 2:**
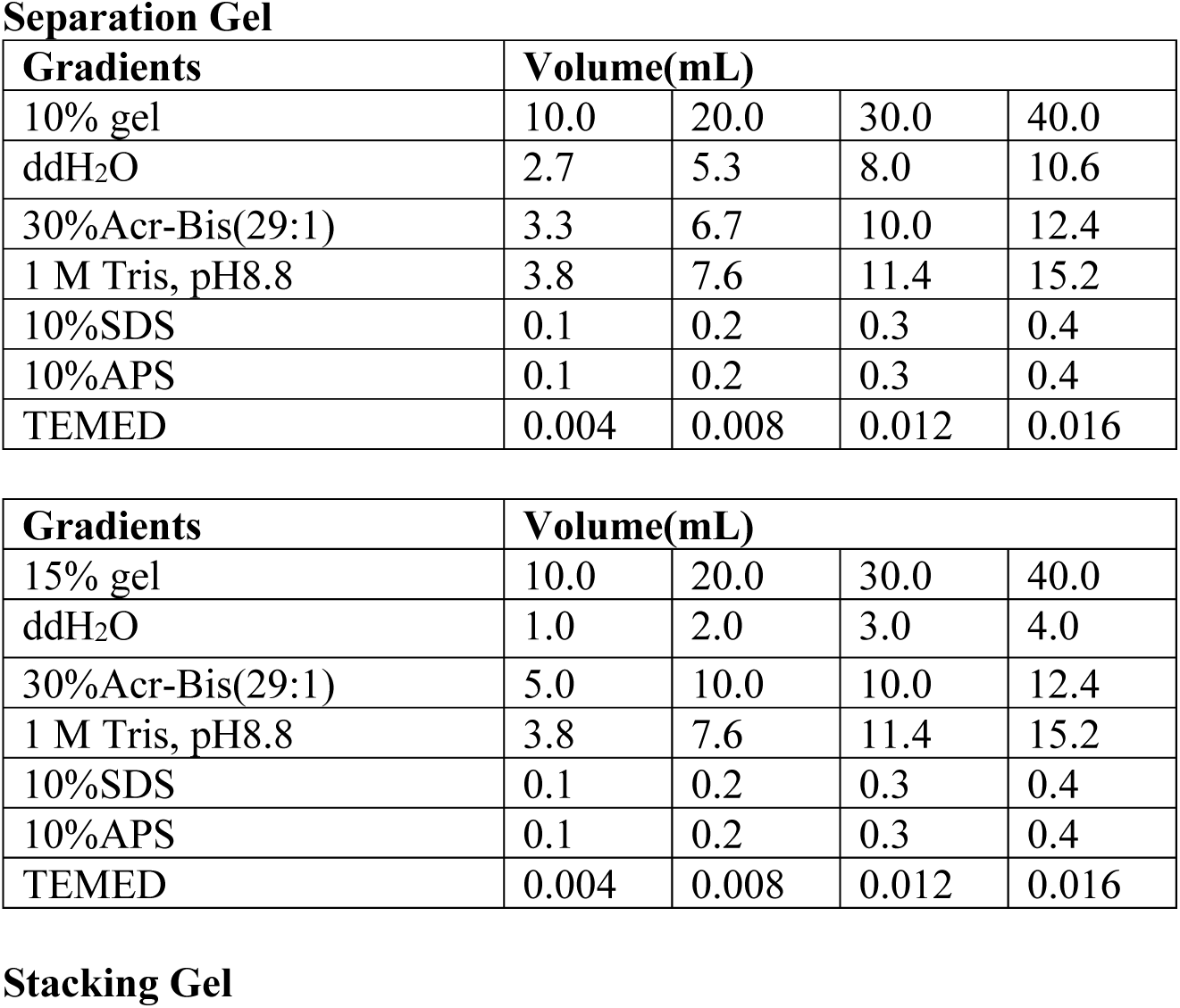
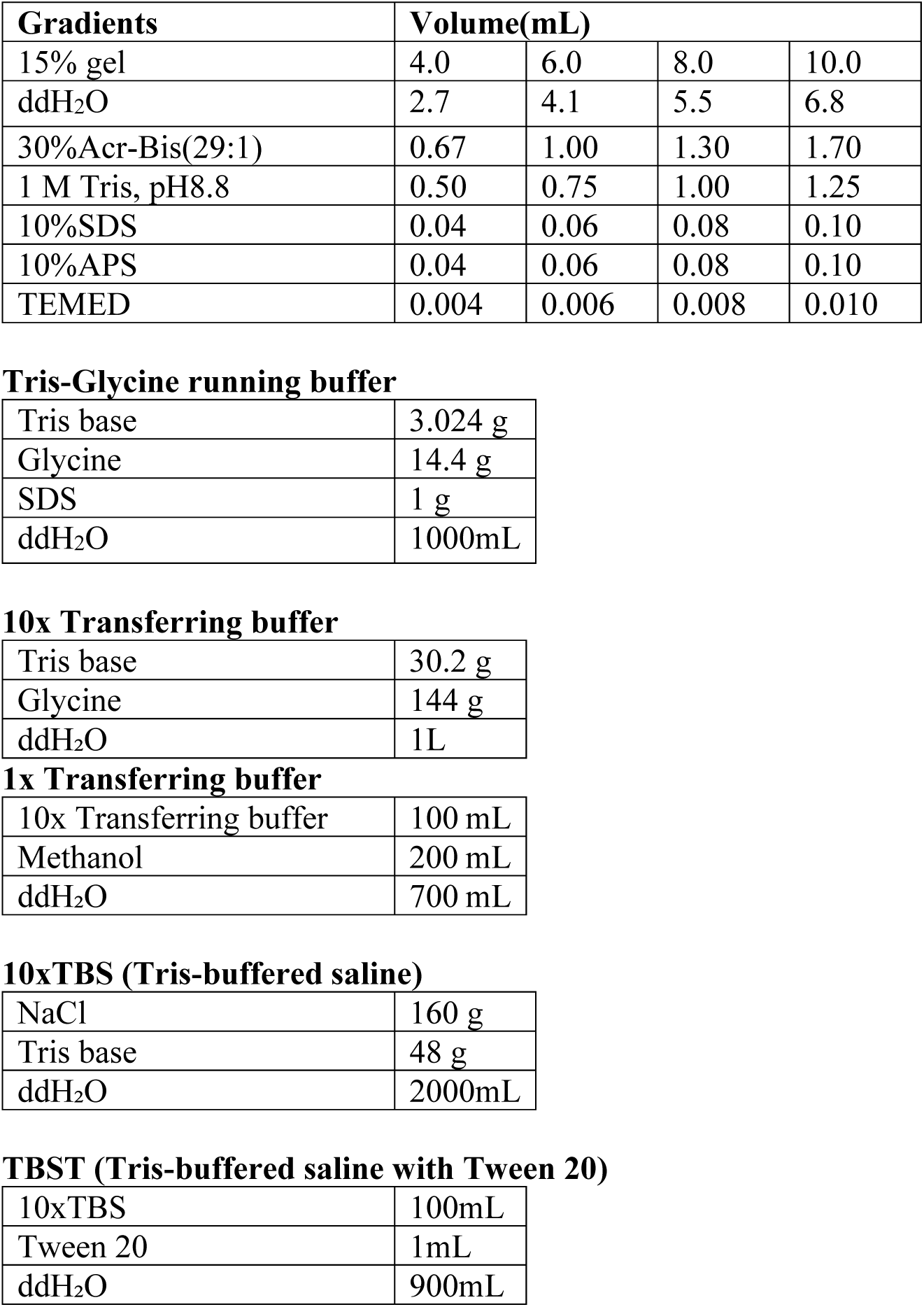
Gel and Buffer Recipes.

| Gradients | Volume(mL) |  |  |  |
| --- | --- | --- | --- | --- |
| 10% gel | 10.0 | 20.0 | 30.0 | 40.0 |
| ddH<sub>2</sub>O | 2.7 | 5.3 | 8.0 | 10.6 |
| 30%Acr-Bis(29:1) | 3.3 | 6.7 | 10.0 | 12.4 |
| 1 M Tris, pH8.8 | 3.8 | 7.6 | 11.4 | 15.2 |
| 10%SDS | 0.1 | 0.2 | 0.3 | 0.4 |
| 10%APS | 0.1 | 0.2 | 0.3 | 0.4 |
| TEMED | 0.004 | 0.008 | 0.012 | 0.016 |
| 15% gel | 10.0 | 20.0 | 30.0 | 40.0 |
| ddH<sub>2</sub>O | 1.0 | 2.0 | 3.0 | 4.0 |
| 30%Acr-Bis(29:1) | 5.0 | 10.0 | 10.0 | 12.4 |
| 1 M Tris, pH8.8 | 3.8 | 7.6 | 11.4 | 15.2 |
| 10%SDS | 0.1 | 0.2 | 0.3 | 0.4 |
| 10%APS | 0.1 | 0.2 | 0.3 | 0.4 |
| TEMED | 0.004 | 0.008 | 0.012 | 0.016 |

Stacking Gel
| **Gradients** | **Volume(mL)** |  |  |  |
| --- | --- | --- | --- | --- |
| 15% gel | 4.0 | 6.0 | 8.0 | 10.0 |
| ddH<sub>2</sub>O | 2.7 | 4.1 | 5.5 | 6.8 |
| 30%Acr-Bis(29:1) | 0.67 | 1.00 | 1.30 | 1.70 |
| 1 M Tris, pH8.8 | 0.50 | 0.75 | 1.00 | 1.25 |
| 10%SDS | 0.04 | 0.06 | 0.08 | 0.10 |
| 10%APS | 0.04 | 0.06 | 0.08 | 0.10 |
| TEMED | 0.004 | 0.006 | 0.008 | 0.010 |

**Tris-Glycine running buffer**
|  |  |
| --- | --- |
| Tris base | 3.024 g |
| Glycine | 14.4 g |
| SDS | 1 g |
| ddH<sub>2</sub>O | 1000mL |

**10x Transferring buffer**
|  |  |
| --- | --- |
| Tris base | 30.2 g |
| Glycine | 144 g |
| ddH<sub>2</sub>O | 1L |

**1x Transferring buffer**
|  |  |
| --- | --- |
| 10x Transferring buffer | 100 mL |
| Methanol | 200 mL |
| ddH<sub>2</sub>O | 700 mL |

**10xTBS (Tris-buffered saline)**
|  |  |
| --- | --- |
| NaCl | 160 g |
| Tris base | 48 g |
| ddH<sub>2</sub>O | 2000mL |

**TBST (Tris-buffered saline with Tween 20)**
|  |  |
| --- | --- |
| 10xTBS | 100mL |
| Tween 20 | 1mL |
| ddH<sub>2</sub>O | 900mL |

**Appendix 3:** Antibody Dilutions.

| Antibody | Dilution |
| --- | --- |
| GAPDH (primary) | 1:1000 |
| Smad2 (primary) | 1:2000 |
| Smad4 (primary) | 1:5000 |
| TGFβ3 (primary) | 1:1000 |
| Goat anti-rabbit (secondary) | 1:5000 |
| Goat anti-mouse (secondary) | 1:5000 |

**Appendix 4:**
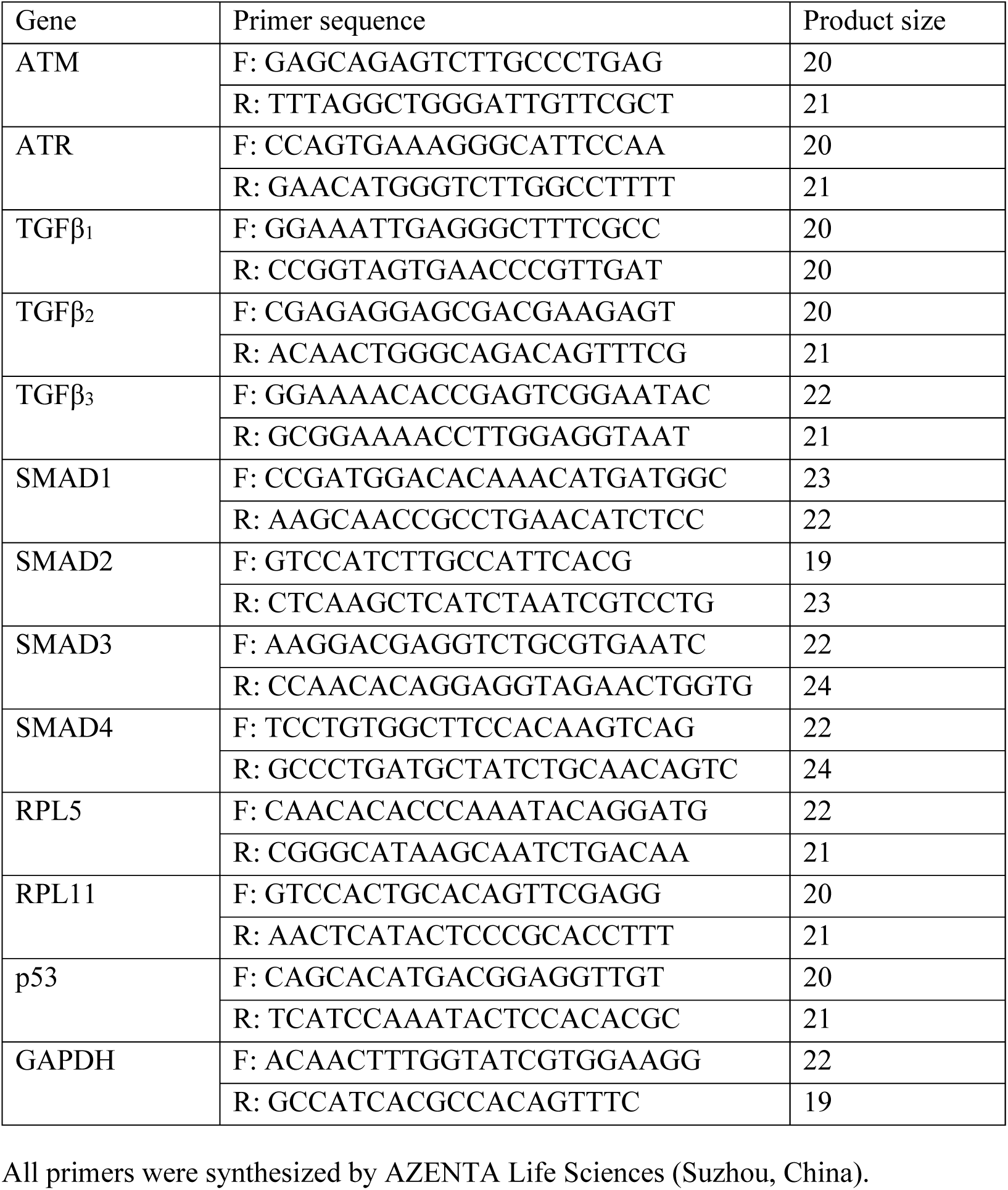
Q-PCR Primers.

## Declarations

### Ethics Approval and Consent to Participate

This study did not include human or animal participants.

### Availability of Data and Material

Data supporting this study are included within the article figures and tables.

### Consent for Publication

All authors read and approved of the manuscript for publication. This study does not include research participants or require their consent.

### Author Contributions

Hongran Ding and Anastasia Tsigkou contributed to the study conception and design. Data collection and analysis were performed by Hongran Ding. Hongran Ding drafted the manuscript. Maya Slack, Holly McClure, Wen Gu, and Helene Gu revised the manuscript. The graphical abstract was designed by Holly McClure. All authors read and approved of the final manuscript. Jason Somarelli, Thomas Schultz, Anastasia Tsigkou, and Ferdinand Kappes supervised this project.

## Acknowledgements

The authors would like to acknowledge Marius Wamsiedel and Chenkai Wu for their supervision of the Global Health Program. The authors would also like to thank Bo Chen for assistance in data collection and analysis, as well as Renata Koviazina and Carly Nabinger for their work in revisions.

